# Protein Synthesis Blockade Prevents Fear Memory Reactivation via Inhibition of Engram Synapse Strengthening

**DOI:** 10.1101/2025.02.05.636389

**Authors:** Ilgang Hong, Yeonjun Kim, Hyunsu Jung, Chang-Ho Kim, Jun-Hyeong Cho, Bong-Kiun Kaang

## Abstract

Memory relies on ensembles of engram cells in the brain. While previous studies have established the existence of these cells, the relationship between cellular and synaptic activity remains unclear. To address this, we applied the dual e-GRASP technique in mice to examine synaptic connectivity between the ventral CA1 and the basal amygdala during memory formation. We found that contextual fear conditioning increased engram-to-engram synapse (engram synapse) density and induced structural potentiation, underscoring their importance in associative memory. Additionally, we investigated the role of protein synthesis in memory formation by inducing retrograde amnesia using anisomycin, a protein synthesis inhibitor. Mice injected with anisomycin once (1xANI) showed impaired natural recall but still displayed fear responses upon optogenetic reactivation of engram cells. In contrast, mice that received four anisomycin injections over six hours (4xANI) showed significantly impaired natural recall and failed to exhibit fear responses upon optogenetic reactivation. This behavioural phenotype was correlated with synaptic changes: 1xANI mice exhibited no significant reduction in engram synapse density but had significantly smaller spine sizes, whereas 4xANI mice showed both a significant reduction in engram synapse density and spine size. Our results indicate that protein synthesis inhibition significantly reduces engram synapse density and spine size, changes that correlate with reductions in fear memory during both natural recall and artificial reactivation.

## Introduction

The concept of the engram refers to the physical trace of a memory, which operates across multiple scales^1^. Engrams have been studied at the synaptic level in various behavioural paradigms—including contextual fear conditioning, auditory fear conditioning, and motor learning—as well as across different memory stages, such as consolidation, extinction, and remote recall^2–5^. However, whether this hypothesis extends beyond the CA3-CA1 circuit in fear learning remains unclear. In this study, we explore the engram synapse hypothesis within the ventral CA1 (vCA1) – basal amygdala (BA) circuit during associative fear learning.

The vCA1 is known to play a critical role in aversive learning, with growing evidence suggesting that engram neurons within this region encode information essential for forming lasting associations such as linking a specific environment to an aversive outcome^6–8^. Conversely, the amygdala is widely recognised for its role in processing emotions, especially fear-related responses^9,10^. Within this framework, activated vCA1 neurons are thought to convey contextual representations to the BA. Although the anatomical and functional connectivity between vCA1 and BA has been demonstrated in both *ex vivo* and *in vivo* studies, direct visualization of this connection at the synaptic level remains largely unexplored.

Protein synthesis is known to be critical in long-lasting forms of synaptic plasticity^11,12^. Applying a protein synthesis inhibitor (PSI) such as cycloheximide or anisomycin in brain slices prior to tetanisation has been shown to reduce the amplitude of field excitatory postsynaptic potentials and impair synaptic transmission during the late phase of long-term potentiation^13–15^. Consequently, inhibiting protein synthesis during acquisition blocks natural recall^16^. As anticipated, the disruption of memory consolidation is accompanied by the loss of enhanced synaptic strength in engram cells^16,17^.

Protein synthesis is essential for natural memory recall, but its role in memory storage and artificial recall remains debated. A single anisomycin injection after memory acquisition impairs natural recall, yet optogenetic reactivation of DG engram cells can still induce memory retrieval^16^. However, when anisomycin is administered alongside tat-Beclin, an autophagy-inducing drug, both natural and artificial recall are disrupted^18^. This raises the question of whether memory storage and artificial recall are critically affected by the magnitude of protein synthesis blockade. Additionally, the synaptic changes associated with protein synthesis blockade remain unclear. Here, we demonstrate that in the vCA1-BA circuit, the impairment of both natural recall and optogenetic reactivation depends on the degree of protein synthesis inhibition, which correlates with reductions in engram synapse density and spine size.

## Results

### Validation of vCA1-BA circuit during contextual fear memory

The inner (stratum radiatum facing) and outer (stratum oriens facing) layers of vCA1 projecting to the BA are known to play distinct roles in anxiogenic and anxiolytic behaviour, but also in fear memory^6,19,20^. To determine which layer projects to the BA, retrograde viral tracing was performed by injecting retrogradely transducing CAV-Cre into the BA of double-floxed tdTomato mice. Retrograde tracing confirmed that BA-projecting vCA1 neurones mainly reside in the outer layer (Fig. S1).

Next, to see whether each neurone can store fear memory as an engram cell, the reactivation ratio between acquisition- and retrieval-activated cells was observed. Acquisition-activated neurones were labelled using c-Fos promoter driven Tet-on system and recall-activated neurones were labelled using c-Fos immunohistochemistry (Fig. S2 A, B). A significant increase in colocalisation ratio was observed in the contextual fear conditioning (CFC) group compared to the naïve group in both vCA1 and BA (Fig. S2 C-E).

### Increased synaptic density and spine size between vCA1 and BA engram cells during memory formation

Next, correlation between associative memory formation and synaptic connectivity between engram cells was assessed. Dual enhanced green fluorescent protein reconstitution across synaptic partners (dual-eGRASP) enables specific labelling of activated dendrites and synapses^2^. Dual-eGRASP utilises split protein reconstitution in the synaptic cleft to label synapses. Using two colours of split protein and fluorescence expression, all connection types between engrams (E) and non-engrams (N) can be fluorescently labelled (Fig. 1A-C, S3A).

**Figure 1.**
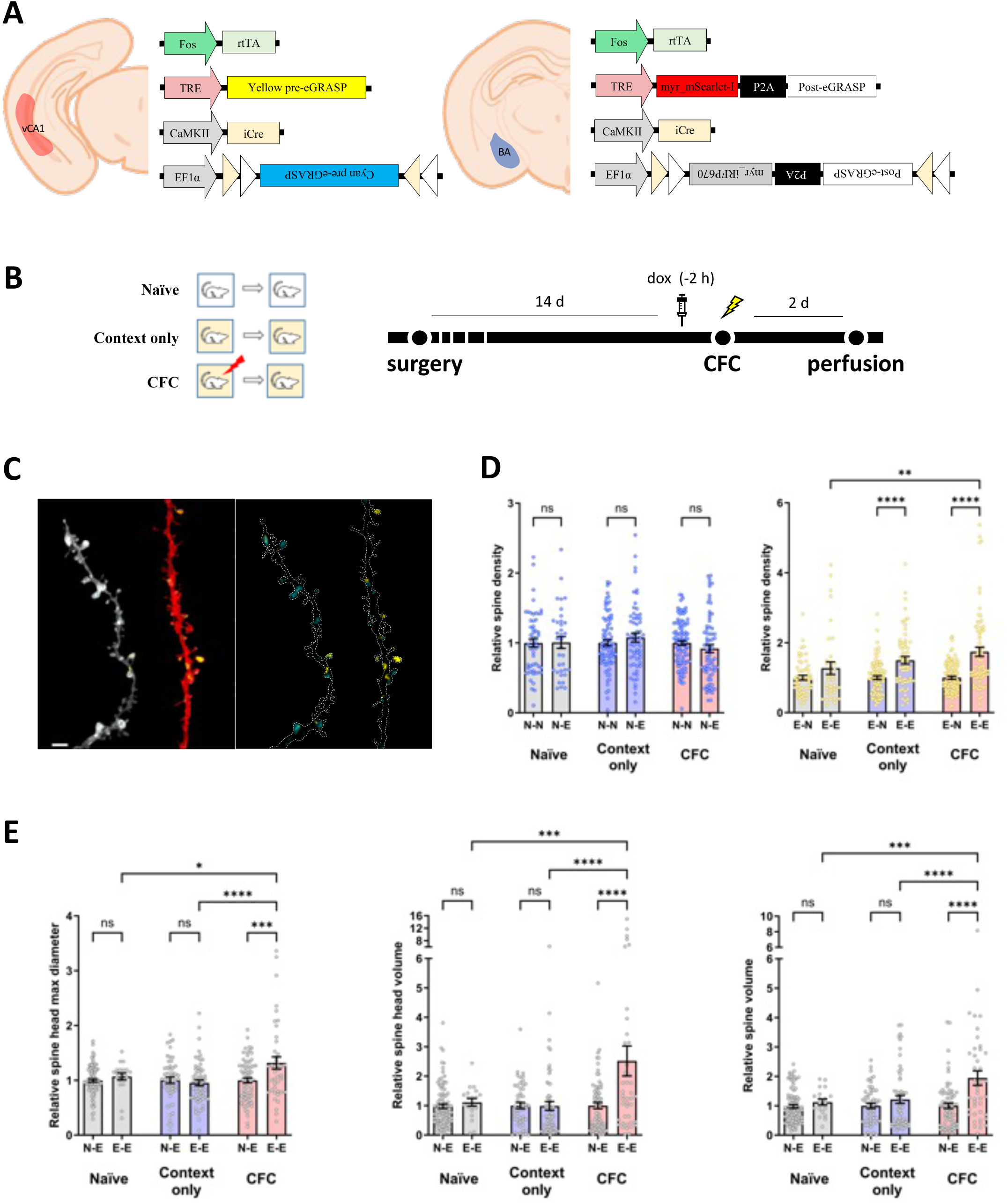
Synaptic density between the vCA1-BA pathway. (**A**) Illustration of virus injection sites. Injection in each site was performed with a cocktail of all the viruses shown for each site. (**B**) Schematic of the experimental protocol. (**C**) Illustrative image depicts both engram (red) and nonengram (gray) dendrites with dual-eGRASP labelling in the BA. Cyan signals represent random vCA1 inputs while yellow signals indicate vCA1 engram inputs. Scale bar : 3 μm (**D**) Normalised cyan/yellow eGRASP per dendritic length. The densities of cyan-only or yellow puncta on mScarlet-I+ dendrites are normalised to the corresponding cyan-only or yellow puncta on iRFP670+ dendrites from the same images to exclude the effect of different number of vCA1 cells expressing each presynaptic component. Each data point represents a dendrite. Naïve group, *n* = 45 for BA non engram dendrites, *n* = 27 for BA engram dendrites; Context only group, *n* = 106 for BA non engram dendrites; *n* = 66 for BA engram dendrites; CFC group, *n* = 73 for BA non engram dendrites; *n* = 62 for BA engram dendrites. Two-way ANOVA followed by Tukey’s multiple comparison test. **P = 0.0063; ****P < 0.0001. Data are presented as mean ± SEM. (**E**) Normalised spine head diameters, spine head volume and spine volume of dendrites from BA engram cells. Naïve group N-E, *n* = 111; E-E, *n* = 32; Context only group N-E, *n* = 107; E-E, *n* = 75; CFC group N-E, *n* = 71; E-E, *n* = 37. Two-way ANOVA followed by Tukey’s multiple comparison test. spine head diameter *P = 0.0315; ***P = 0.0001, spine head volume ***P = 0.0004; ****P < 0.0001, spine volume ***P = 0.0001; ****P < 0.0001. Data are presented as mean ± SEM.

Using this dual-eGRASP technique, synaptic strengthening was quantified in the vCA1-BA circuit after CFC (Fig. 1B, S3B). No significant difference in synaptic density was observed between N-N and N-E across all experimental groups while significantly higher density of E-E synapses was observed in the CFC group compared to the E-N synapses (Fig. 1D). This suggests recruitment of synapses between engram neurones during acquisition of associative memory. Subsequently, morphological changes were investigated to confirm structural plasticity in the spines during associative memory formation. Morphological analysis revealed a significant increase in spine head diameter, spine head volume, and spine volume of E-E synapses in the CFC group compared to N-E synapses (Fig. 1E). The context only group exhibited no significant differences in spine head size of E-E synapses. We conclude that fear conditioning induced structural plasticity specifically in E-E synapses. No significant difference was found in any of the four metrics of structural plasticity between N-N and N-E synapse in the three behavioural groups (Fig. S3F). Consequently, these results indicate that engram synapses undergo structural plasticity during associative fear learning in the vCA1-BA circuit.

### *De novo* protein quantification using NCAA labelling

A key aspect of memory consolidation theory suggests that *de novo* protein synthesis is essential for synaptic plasticity^21,22^. The use of PSIs has been crucial in advancing this theory, as numerous studies have demonstrated that administering PSIs during memory acquisition impairs natural recall in mice^16^. Therefore, as an indirect method to inhibit synaptic changes observed in Fig 1, we established behavioral conditions for PSI treatment with greater specificity.

First, to establish experimental conditions for extensive protein synthesis inhibition, noncanonical amino acids (NCAA) labelling using biorthogonal noncanonical amino acid tagging (BONCAT)^23–25^ was performed as mice were injected with PSI. Given that the reported time window for memory consolidation extends up to 6 hours after the event^26^, initial approach aimed to extend the duration of the PSI treatment^27–29^. It has been previously reported using an electrophysiological approach that four consecutive injections of anisomycin weaken E-E synaptic strengthening^7^. Therefore, mice received four consecutive anisomycin injections (4xANI), or a single anisomycin injection followed by three consecutive saline injections (1xANI), or 4 consecutive saline injections (SAL). Immediately after the CFC, mice were intraperitoneally injected with L-methionine analog L-azidohomoalanine (AHA) which tags newly synthesizing proteins and brains were snap frozen 16 hours after CFC (Fig. 2A). Quantification of NCAA labelled proteins by western blotting showed that 4xANI mice displayed protein intensity comparable to that of AHA(-) mice (Fig. 2B, C). These results suggest that 4xANI injections effectively suppresses protein synthesis.

**Fig 2.**
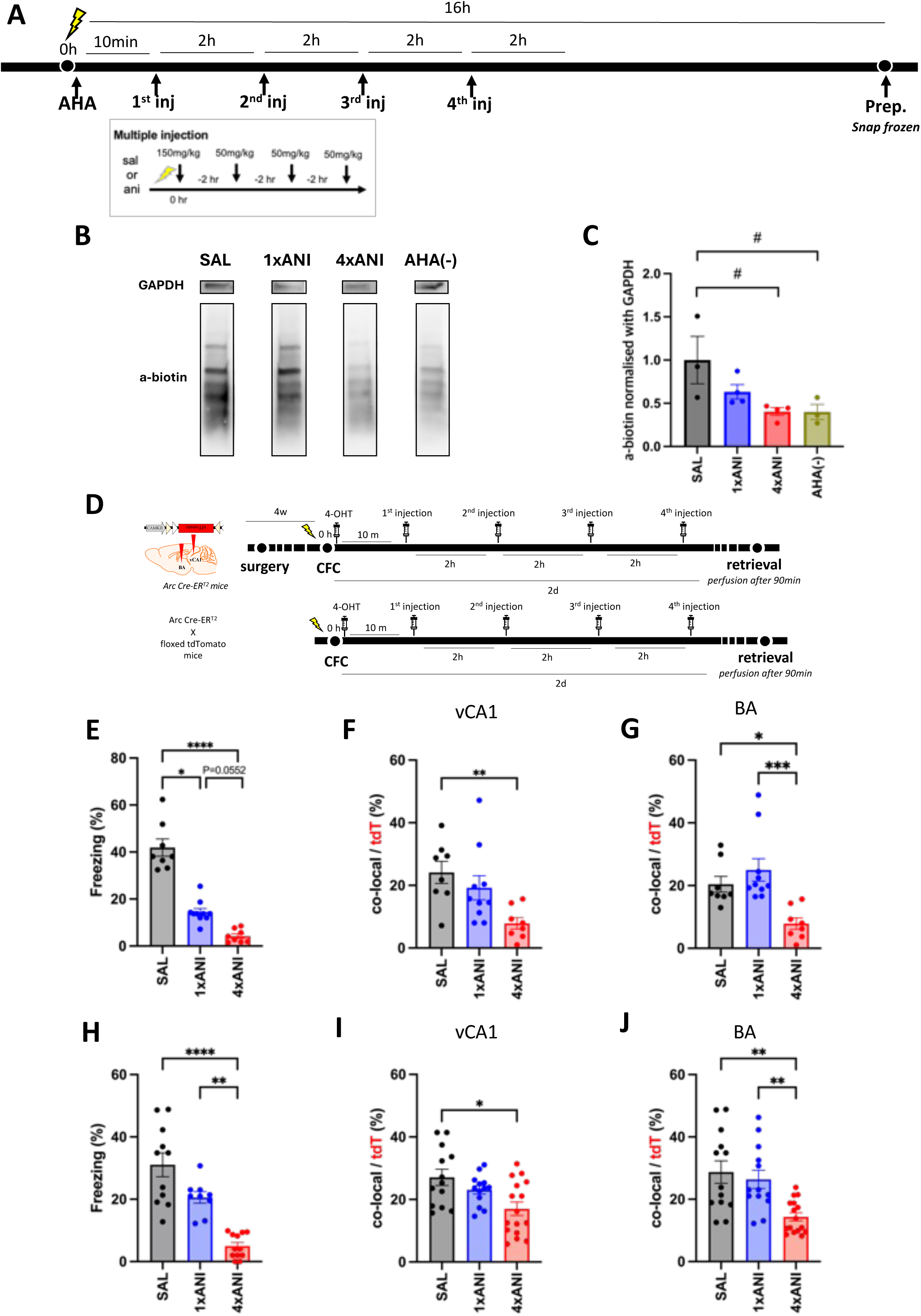
Engram cells are less reactivated by fear recall with multiple anisomycin injections. (A) Behavioural procedure for NCAA labelling. (B) Western blot image to tag *de novo* synthesized proteins during memory acquisition (C) Quantification of the de novo protein synthesis level normalised using GAPDH. SAL group, N = 3; 1xANI group, N = 4; 4xANI group, N = 4; AHA(-) group, N = 3. Kruskal-Wallis test followed by two-stage linear step-up procedure of Bejamini, Kriegar and Yekutieli. #P = 0.0776. Data are presented as mean ± SEM. (D) Behavioural procedure for testing engram reactivation ratio. (E-J) Data of engram reactivation analysis for Arc-CreERT2 mice expressing viral CaMKII-DIO-tdTomato (E-G) or crossed with floxed tdTomato (H-J). (E, H) Freezing level during memory recall session. 4xANI group showed lower freezing level compared to that of 1xANI group in both conditions. Engram reactivation ratio of vCA1 neurones (F, I) and BA neurones (G, J). Significantly lower reactivation ratio of BA neurones was observed in 4xANI group compared to 1xANI group. (E-G) SAL group, N = 8; 1xANI group, N = 10; 4xANI group, N = 8. (H-J) SAL group, N = 13; 1xANI group, N = 13; 4xANI group, N = 16. Kruskal-Wallis test followed by Dunn’s multiple comparison test. (E) *P = 0.0336; ****P < 0.0001. (F) **P = 0.0045. (G) *P = 0.0148; ***P = 0.0003. (H) ****P < 0.0001; **P = 0.0033. (I) *P = 0.0230. (J) SAL vs. 4xANI, **P = 0.0027; 1xANI vs. 4xANI, **P = 0.0041. Data are presented as mean ± SEM.

### Multiple injections and cocktail of drugs induce retrograde amnesia more effectively

Building on NCAA studies, robust protein synthesis inhibition conditions were applied *in vivo*. For multiple injections, four consecutive injections of saline or anisomycin were delivered (Fig. 2A, D). In another experimental condition, a stronger inhibition was achieved with a single injection by using cycloheximide along with anisomycin to inhibit two different ribosomal sites (Fig. S5). In line with the previous study, a single injection of anisomycin decreased natural recall of fear memory^16^, while multiple injections of anisomycin and the cocktail group (CKT) exhibited significantly lower freezing levels compared to the single injected group (Fig. 2E, 2H, S5A-B). Therefore, these data confirm that extensive protein synthesis inhibition by multiple injections of anisomycin or a cocktail of protein synthesis inhibitors significantly impairs natural recall of fear memory.

To test whether the memory impairment by injection of PSI leads to impaired engram reactivation, Arc-CreERT2 crossed with floxed tdTomato mice were fear conditioned followed by 4-OHT and a series of PSI injections (Fig. 2D). Immunohistochemistry analysis showed no significant difference in the ratio of tdTomato^+^ or c-Fos^+^ neurones in all three groups (Fig. S4).

However, the reactivation ratio between activated neurones during the conditioning and memory recall was significantly lower in the 4xANI group in both vCA1 and BA (Fig. 2I, J). Significant differences in reactivation ratio between groups were replicated using Arc-CreERT2 mice expressing CaMKII-DIO-tdTomato (Fig. 2F, G). Moreover, the single injection of the cocktail significantly reduced the reactivation ratio in the BA compared to treatment with anisomycin alone (Fig. S5). Taken together, these data suggest that multiple injections of anisomycin significantly disrupt the reactivation of engram cells and impair memory recall.

We next examined whether amnesic memory can be recalled using optogenetics. To reactivate neurones specifically activated during the CFC under different PSI treatments, EF1a-DIO-ChR2-eYFP was bilaterally expressed in the vCA1 of Arc-CreERT2 mice (Fig. 3B, S6A). To reach sufficient virus expression in axon terminals in the BA, mice were tested five days after the CFC (Fig. S6B, C). Restoration of memory from amnesia has been observed by directly stimulating anisomycin-treated DG engram cells with light; this preferential, protein synthesis-independent functional engram connectivity was observed even eight days after the initial training^30^. All groups of mice reported here showed robust freezing acquisition (Fig. 3A). Five days after training, mice were introduced to a novel environment for blue light stimulation. In line with prior research, neither group displayed freezing behaviour when the blue light was off, but SAL groups and 1xANI groups exhibited a significant increase in freezing behaviour when the blue light was on (Fig. 3C-D, S7C-D). Notably, in the CKT and 4xANI group, mice did not display any freezing level during the light on and off sessions (Fig. 3E, Fig. S7E). To assess the conditioned response to recall cues, mice were once more evaluated in the fear-conditioned chamber 24 hours after the optical stimulation. Consistent with our previous data, SAL mice showed robust freezing while 4xANI showed minimal freezing behaviour (Fig. 3B).

**Figure 3.**
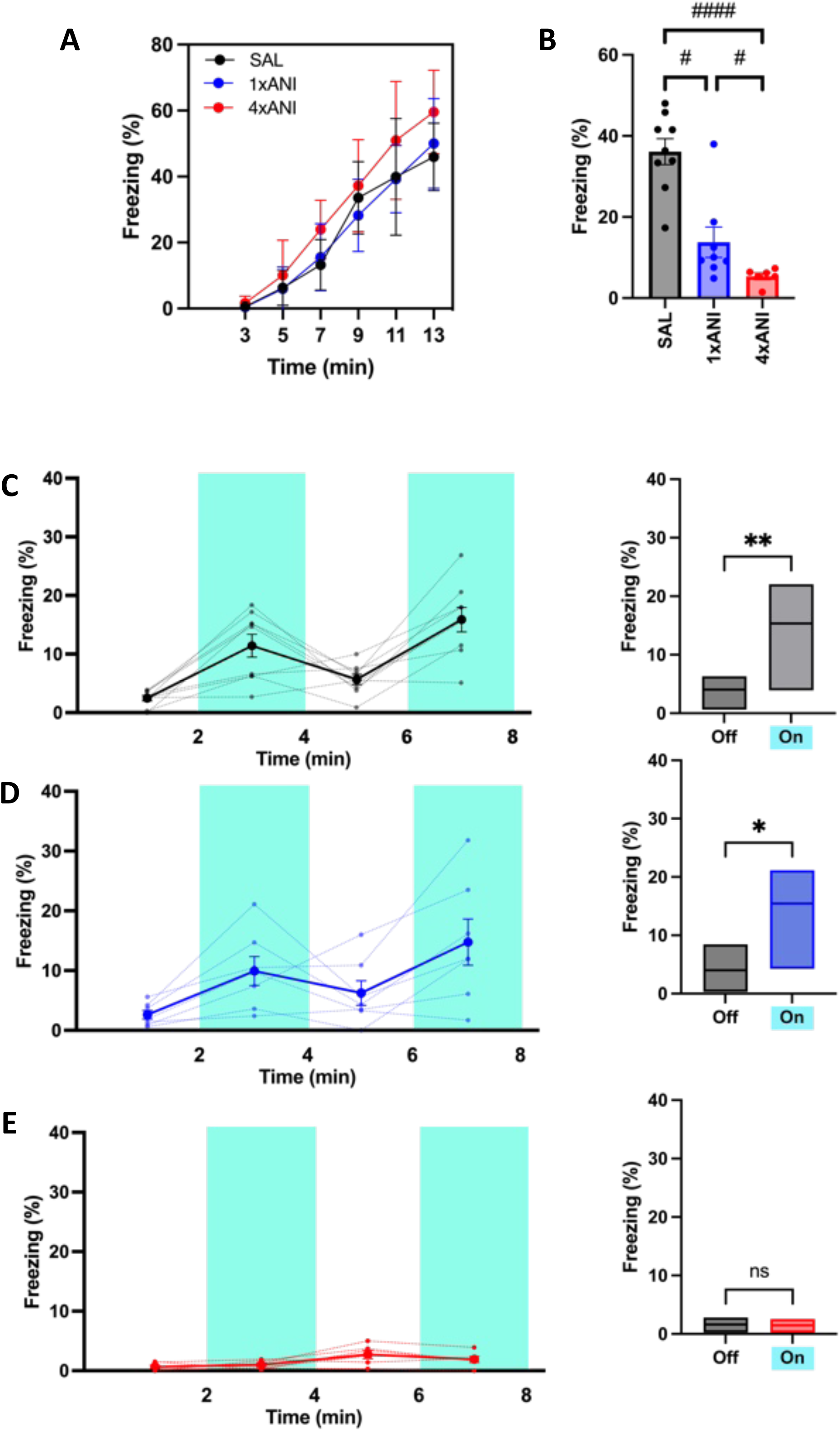
Memory cannot be artificially recalled when protein synthesis was blocked for extensive time during memory acquisition. (A) Freezing level during memory acquisition. Data are presented as mean ± SEM. (B) Freezing level during contextual fear memory retrieval. SAL group, N = 9; 1xANI group, N = 7; 4xANI group, N = 6. Kruskal-Wallis test followed by two-stage linear step-up procedure of Bejamini, Kriegar and Yekutieli. SAL vs. 1xANI, #q = 0.0193; 1xANI vs. 4xANI, #q = 0.0859; ####q < 0.0001. Data are presented as mean ± SEM. (C-E) Optogenetic activation of vCA1 engram axon terminals in BA of mice treated with PSI during memory acquisition. Mice expressing ChR2 in vCA1 engram cells from CFC show greater freezing during light-on in saline treated control group (C) or by a single treatment of anisomycin (D) while absence of freezing behaviour was shown in mice treated with multiple injection of anisomycin (E). Wilcoxon test. SAL group, **P = 0.0078; 1xANI group, *P = 0.0156. Data are presented as mean ± SEM (left) or median (right).

The saline group displayed robust freezing behaviour, whereas the anisomycin group with single injection resulted in significantly reduced freezing behaviour during cued recall (Fig. S7B). The 4xANI and CKT groups exhibited a freezing level similar to naïve mice. These data indicate that the connectivity of vCA1 engram cells to the downstream BA is enhanced during memory acquisition only if the protein synthesis blockade is not extensive, while an extensive blockade inhibits synaptic strengthening between the regions, thus impairing memory. Collectively, these data provide evidence supporting the critical role of engram synapses in associative memory by indirectly modulating engram synapse potentiation by multiple PSI injections.

To investigate the synaptic changes associated with memory under extensive protein synthesis blockade, we applied the dual-eGRASP technique (Fig. 1A). Mice underwent CFC followed by serial injections of either anisomycin or saline (Fig. 4A). Consistent with our previous findings, the group that received four injections of anisomycin (4xANI) exhibited significantly impaired fear memory retrieval compared to the SAL group (Figs. 3B and 4B). Dual-eGRASP analysis revealed an increased density of E-E synapses across all behavioural groups, while no significant differences were observed in N-E synapse density (Figs. 1D and 4C). However, a notable reduction in E-E synapse density was observed in the 4xANI group compared to the 1xANI, with no significant difference between the SAL and 1xANI groups (Fig. 4C). This reduction in E-E synapse density may explain the failure of optogenetic reactivation observed in the 4xANI group (Fig. 3E). Additionally, spine morphology analysis revealed increases in spine head diameter, spine head volume, and overall spine volume in the E-E synapses of the SAL group (Figs. 1E and 4D, E). Interestingly, these metrics showed no significant changes in the 1xANI or 4xANI groups, with spine dimensions in the 1xANI group being significantly smaller than those in the SAL group. These findings suggest that the reduction in spine dimensions correlates with impaired natural memory recall observed in the 1xANI group (Fig. 3D).

**Figure 4.**
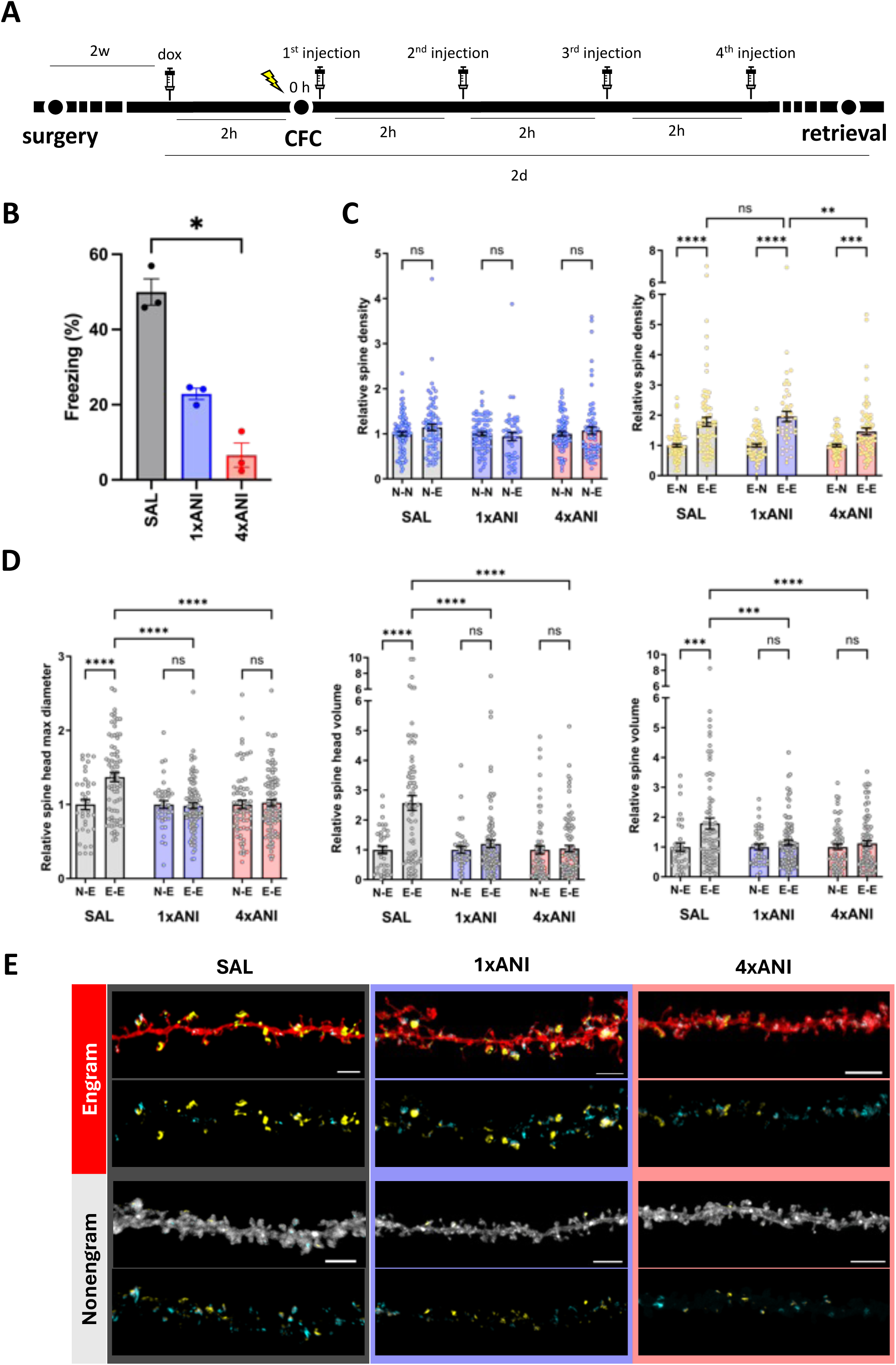
Reduced E-E Synapse Density using dual-eGRASP correlates with impaired optogenetic reactivation in 4xANI. (A) Schematic of the behavioural experiment. Mice were intraperitoneally injected with doxycycline 2 hours prior to CFC. After CFC, 4 consecutive anisomycin was intraperitoneally delivered. 2 days following CFC, mice were subjected to fear memory retrieval. (B) Freezing level during contextual fear memory retrieval for the dual-eGRASP experiment. SAL group, N = 3; 1xANI group, N = 3; 4xANI group, N = 3. Kruskal-Wallis test followed by Dunn’s multiple comparison test. *P = 0.0219. Data are presented as mean ± SEM. (C) Normalised cyan/yellow eGRASP per dendritic length. The densities of cyan-only or yellow puncta on red dendrites are normalised to the corresponding cyan-only or yellow puncta on near-infrared dendrites from the same images to exclude the effect of different number of vCA1 cells expressing each presynaptic component. Each data point represents a dendrite. SAL group, *n* = 85 for BA nonengram dendrites, *n* = 74 for BA engram dendrites; 1xANI group, *n* = 70 for BA nonengram dendrites; *n* = 45 for BA engram dendrites; 4xANI group, *n* = 75 for BA nonengram dendrites; *n* = 69 for BA engram dendrites. Two-way ANOVA followed by Tukey’s multiple comparison test. **P = 0.0043; ***P = 0.0008; ****P < 0.0001. Data are presented as mean ± SEM. (D) Normalised spine head diameter, spine head volume and spine volume of dendrites from BA engram cells. SAL group N-E, *n* = 36; E-E, *n* = 76; 1xANI group N-E, *n* = 39; E-E, *n* = 93; 4xANI group N-E, *n* = 66; E-E, *n* = 93. Two-way ANOVA followed by Turkey’s multiple comparison test. spine head diameter ****P < 0.0001, spine head volume ****P < 0.0001, spine volume ***P = 0.0001; ****P < 0.0001. Data are presented as mean ± SEM. (E) Representative confocal image of both engram and nonengram dendrites from all three behavioural groups. Scale bar : SAL engram, 3 μm; SAL nonengram, 4 μm; 1xANI engram, 4 μm; 1xANI nonengram, 5 μm; 4xANI engram, 5 μm; 4xANI nonengram, 6 μm.

## Discussion

It has been challenging to determine whether memory formation strengthens the synapses between engram cells in interconnected brain regions^31^. This challenge stems from the difficulty in distinguishing presynaptic regions formed by engram cells from those formed by non-engram cells. The modification of the GRASP technique has provided the solution enabling the visualisation and analysis of spine density and morphology^2^. Additionally, the position of the colour-determining domain in the presynaptic neurone (cyan/yellow pre-eGRASP) and the common domain in the postsynaptic neurone (post-eGRASP), allowed us to visualize two synaptic populations originating from two functionally distinct presynaptic neuronal populations: one involved in encoding the engram and the other not, both projecting onto a single postsynaptic neurone^2^.

Using the dual-eGRASP technique, we investigate the hypothesis that contextual fear learning is associated with the enhancement of synapses between engram cells, the so-called synaptic engram hypothesis^32^. Engram synapses have been shown to be critical in memory acquisition and extinction where changes in engram synapse density and spine head size were observed^2,3^. Our results showed similar lines of data, where increased synapse density, spine head size, spine volume, and spine head volume were observed specifically in E-E synapses of vCA1-BA circuit (Fig. 1D, E). Therefore, these data suggest that the engram synapse hypothesis can also be applied to the vCA1-BA circuit during associative fear learning.

Along with the CFC group, the context only group also exhibited an increase in E-E synaptic density (Fig. 1D). This data is consistent with our previous findings within the CA3-CA1 pathway, where the context only group demonstrated significant synaptic changes^2^. This increase in synaptic density may be attributed to Hebbian plasticity between vCA1 engram neurones and BA neurones activated by emotions other than fear, such as anxiety, generated during exposure to the novel context^33^ as BA is also known to mediate anxiogenic behaviour^20^. Unlike synaptic activity of the vCA1 region drives the increase in synaptic density, although it is not sufficiently pronounced to increase spine size.

Moreover, to validate the hypothesis that modification of E-E synaptic connectivity is necessary for memory acquisition, we administered PSI to induce retrograde amnesia. In this study, we set optimal conditions for inducing extensive retrograde amnesia by extending the time window of protein synthesis inhibition or simultaneously targeting two ribosomal sites. These experimental conditions induce a more pronounced effectiveness in impairing memory recall.

It has been reported that a memory that is impaired by inhibiting synaptic potentiation using PSI can later be artificially reactivated^16^. Another study has demonstrated that inhibiting protein synthesis, coupled with the induction of autophagy, results in irretrievable memory^18^. To assess whether varying degrees of protein synthesis inhibition produce different behavioural outcomes, we tested multiple groups to identify the optimal condition for inducing memory impairment. This approach led to the discovery that when mice undergo an extensive inhibition of protein synthesis, optogenetic reactivation of engram cells can no longer induce memory recall. Multiple injections of anisomycin showed no significant changes to the number of engram cells (Fig. S4D). Also, E-E synapses showed increased density and enlarged structural morphology during associative fear learning (Fig. 1D, E). Therefore, we propose that the failure of optogenetic reactivation in mice subjected to multiple anisomycin injections is attributed to a reduction in E-E synaptic density caused by extensive protein synthesis inhibition (Fig. 4B). On the other hand, we suggest that the impaired natural recall observed in both 1xANI and 4xANI groups is driven by the lack of engram synapse enlargement (Fig.4C). Numerous studies have provided evidence that neither single nor multiple injections of anisomycin are sufficient to block synaptic plasticity to a level comparable to the naïve group^7,16^. Although the AMPAR/NMDAR ratio was lower, EPSC frequency remained elevated, suggesting that synaptic strength weakened while E-E synaptic connections increased^7^. We can conclude that our experimental conditions were sufficient to impair memory recall in both natural and optogenetic reactivation contexts (Fig. 3, 4). However, the residual freezing behaviour and the observed increase in engram synapse density needs further investigation which may be regulated by protein synthesis independent mechanisms such as remodelling of actin skeleton.

Limitations of the current study arise from the fact that anisomycin affects protein synthesis in a non-discriminatory way. Anisomycin was given by intraperitoneal injection, thereby affecting the whole mouse body. To rule out the possibility that protein synthesis blockade from other regions has contributed to the failure of memory storage, future studies should block protein synthesis in a brain region-specific and engram-specific manner, possibly through the use of genetically encodable PSI^34^.

## Materials and methods

### Animals

All experiments were conducted in accordance with the regulations and guidelines of the Institutional Animal Care and Use Committees (IACUC) of Seoul National University. Investigators were not blinded to the genotypes of the mice.

Wild-type C57BL/6N mice were used for dual-eGRASP experiments, while wild-type C57BL/6J mice were used for BONCAT. Heterozygous Arc-CreERT2 (Jackson Laboratory Stock # 022357) mice, obtained by crossing wild-type C57BL6/J and Arc-CreERT2 (+/−) mice, were utilised to label engram neurones and for optogenetic experiments. Heterozygous Arc-CreERT2; Ai14 mice, obtained by crossing Ai14 (Jackson Laboratory Stock # 007914) and Arc-CreERT2 (+/−) mice, were used to label engram neurones in the experiments.

Mice were housed in standard laboratory cages with disposable bedding on a standard light cycle (09h-21h). Mice had access to food and water *ad libitum* and were socially housed in numbers of three to five littermates until surgery. Behavioural experiments were performed during the light cycle. Male and/or female mice were utilised in the experiments with details provided in the figures or figure legends

### AAV production

Adeno-associated virus serotype 1/2 (AAV1/2; AAV particle containing both serotype 1 and 2 capsids) was used for injection as previously described^2^.

The procedure for packaging plasmids into adeno-associated virus serotype 9 (AAV9) involved the following steps: plasmids containing the construct of interest flanked by AAV2 ITR, pAd-ΔF6, and pRep/Cap9 were co-transfected into the AAVpro 293T Cell Line (TAKARA, cat# 632273). The transfected cells were then incubated in DMEM with 10% v/v FBS for five days in a 150 mm culture dish. On the day of harvest, culture medium was collected and the remaining cells were lysed to extract any virus they contained. After centrifugation at 13,490 g for 60 minutes at 4°C, the supernatants were carefully loaded into ultracentrifugation tubes filled with iodixanol gradient solutions.

AAV was purified from the cell lysate using iodixanol density-gradient ultracentrifugation as previously described^35^. 40% iodixanol solutions were withdrawn using 5 mL syringes and further concentrated using Amicon Ultra 100-kDa MWCO ultrafiltration centrifugal devices (Merck, cat# UFC910024).

### Virus-mediated gene expression

For c-Fos labelling experiments, the virus comprised AAV1/2-Fos-rtTA mixed with AAV1/2-TRE-mCherryNuc. Virus was bidirectionally injected into both vCA1 (500 nl; AP: -3.1 mm, ML: ±3.4 mm DV: -3.8 mm) and BA (500 nl; AP: -1.5 mm ML: ±3.0 mm, DV: -4.9 mm).

AAV9-EF1α-DIO-ChR2-EYFP was acquired from Institute for Basic Science Recombinant Virus Packaging Facility.

### Stereotaxic surgery procedure

Mice were anaesthetised by intraperitoneal injection of 25 mg·kg^-1^ ketamine/xylazine and positioned on a stereotaxic apparatus (Stoelting Co.). Ophthalmic ointment was applied to the eyes as a preventive measure against dryness. Bilateral craniotomies were performed using a 0.5 mm diameter drill and the viruses were injected using a micro-needle attached to a 10 mL Hamilton micro syringe (Hamilton; cat# 701LT). A syringe driver (WPI, cat# SP3101-PLUS) was used to maintain the speed of the injection at 0.125 ul·min^-1^. The needle was slowly lowered to the target site and remained for 2 minutes before beginning the injection. After injection, the needle stayed for 10 minutes before it was slowly withdrawn. The incision was sutured, and mice were allowed to recover for 2 weeks before the start of behavioural experiment.

For the optogenetic experiment, a custom implant containing two optic fibres (Newdoon; 0.37 NA 1.25 mm core diameter) was lowered above the injection site at the following coordinate relative to bregma (mm): AP: -1.5 ML: ±3.0 DV: -4.0 for BA. A jewellery screw was screwed into the skull near the bregma. Mice were allowed to recover for at least 2 weeks before behavioural experiments.

### Contextual fear conditioning

Two weeks following AAV injection, all mice underwent a conditioning procedure. Prior to conditioning, each mouse was housed individually for eight days and was subjected to daily handling sessions lasting five minutes for seven consecutive days. Mice were randomly assigned to either the CFC group or the context only control group. Mice were transferred from the vivarium to a holding room adjacent to the testing area at least 30 minutes prior to the start of the experiment. Furthermore, each mouse was transported individually to the conditioning room.

The conditioning chamber measuring 7”W × 7”D × 12”H (Coulbourn Instruments; model H10-11M-TC) was used. The chamber featured two aluminium side walls and a ceiling with an acrylic door. It was situated within a larger sound-attenuating enclosure. Throughout the experimental procedures, a white overhead light continuously illuminated the conditioning chamber. The chamber itself was equipped with a straight stainless-steel rod floor which was cleaned using a 70% ethanol solution and distilled water between each use.

Two hours prior to the conditioning procedure, an intraperitoneal injection of 250 μL of a 5 mg·ml^-1^ Doxycycline solution dissolved in saline was administered. For contextual fear conditioning, mice were exposed to the context for 180 seconds, after which five-foot shocks were delivered where each foot shock lasted 2 seconds with an interval of 2 minutes. Following the conditioning session, mice were returned to their respective home cages. During the retrieval test, the percentage of time spent freezing was calculated and averaged across the entire 3-minute retrieval session.

### BONCAT

General procedures for BONCAT was followed as previously described^23–25^. AHA was dissolved in PBS (NaCl 137 mM, KCl 2.7 mM, Na2HPO4 4.3 mM, KH2PO4 1.47 mM) and administered via intraperitoneal injection. The labelling duration was set as 16 hours following the established treatment protocol for optimal results^36^ with a dosage of 100 μg NCAA·gbw^-1^ (gram body weight). After the designated treatment period, mice were deeply anesthetised with isoflurane followed by brain removal. The vCA1 and BA were sliced in the artificial cerebrospinal fluid (NaCl 124 mM, KCl 3 mM, NaHCO3 26 mM, NaH2PO4 1.25 mM, MgSO4 2 mM, D-glucose 15 mM, CaCl2 2 mM) carbonated with 95% O2 and 5% CO2 into 300μm sections using a vibratome (Leica; VT1200S). Brains were dissected under a dissecting microscope and frozen using liquid nitrogen.

To create working stocks of 4-Hydroxytamoxifen (4-OHT) at a concentration of 20 mg·ml^-1^, 50 mg of 4-OHT (Sigma Aldrich, cat# H6278) were dissolved in 2.5 ml of 100% ethanol. These working stocks of 4-OHT were stored at -20°C for a maximum of three months. On the day of use, working stocks were further diluted with a mixture of castor oil and sunflower oil at a 1:4 ratio (Sigma Aldrich, cat# 259853 and cat# S5007) via vacuum centrifugation resulting in a final concentration of 10 mg·ml-1.

Anisomycin (Tocris, cat# 1290) and cycloheximide (Sigma Aldrich, cat# 239763-M) were stored at -20°C up to a year, and the powder was dissolved with saline on the day of use.

### BONCAT purification and Immunoblotting

General procedures for BONCAT was followed as previously described^23–25^. The protein concentration was assessed using a Pierce BCA protein assay kit (Thermo Fisher Scientific) following the manufacturer’s guidelines. Subsequently, an equal volume of protein samples was subjected to electrophoresis on a 4 to 12% gradient gel (BIO-RAD, cat#4561084) and transferred onto a PVDF membrane (Invitrogen, cat#IB24001). The membranes were blocked using 4% bovine serum albumin in Tris-Buffered Saline, 0.1% Tween® 20 (TBST) and then incubated with anti-Biotin (1:1000, cat#5597, Cell Signalling) overnight at 4°C. Blots were washed three times in TBST and then incubated with the secondary antibody (anti-rabbit HRP, 1:5000, Invitrogen; cat#Q-11071MP) at room temperature for 2 hours. The bands were visualised utilising a West-Q Chemiluminescent Substrate Kit (GenDEPOT, W3651-012) and the ChemiDoc XRS+ system (Bio-Rad; cat#721BR02655). Signal detection and quantification were performed using the ChemiDoc MP device (Bio-Rad).

### Optogenetics

For the single injection groups, mice were either injected with saline, anisomycin, or anisomycin + cycloheximide right after conditioning. For the multiple injection groups, mice were first injected with saline or anisomycin, followed by three injections with saline or anisomycin at 2 hours of intervals. 4-OHT was injected intraperitoneally for 50 mg·kg^-1^10 minutes after the first injection. For the immunohistochemistry, mice were retrieved two days after the conditioning and perfused 90 minutes after retrieval. For optogenetics, mice were habituated to optical cables for at least 30 minutes before the experiment. From 30 to 60 minutes of habituation, mice were placed in the acrylic chamber. Mice were allowed to freely move within the acrylic chamber and freezing behaviour was recorded for 8 minutes, with 2 minutes of two consecutive lights ‘off’ and ‘on’ sessions.

### Immunohistochemistry

90 minutes after retrieval, mice were deeply anesthetised using ketamine/xylazine (15 mg·kg^-^ 1) followed by transcranial perfusion with 15 ml of PBS and 15 ml of 4% PFA. Brains were removed, post-fixed overnight at 4°C in 4% PFA and dehydrated in a 30% sucrose solution in PBS at 4°C for two days. The brain was rinsed with PBS and subsequently frozen at -80°C for a minimum of one hour. Coronal sectioning was performed using a cryostat (Leica) with a thickness of 30 μm. The sections were rinsed with 1 mL of PBS for 5 minutes each time at room temperature (RT), 120 rpm. Then 500 μL of a blocking solution (5% normal goat serum, 0.3% triton X-100 in PBS) was applied to each sample for 1hr RT, 80 rpm. The primary c-Fos antibody (SySy, cat# 226 003) was applied at a concentration of 1:1000 for 24 hours on a shaker at 4 °C. Sections were washed three times with PBS for 5 minutes each time at room temperature (RT), 120 rpm. A secondary antibody (Alexa-488; 1:500) was applied at a concentration of 1:500 for 2 hours at 80 rpm. Finally, sections were washed with 1 mL of PBS three times for 5 minutes each time at 120 rpm, RT. The second washing step included DAPI staining at a concentration of 1:1000 for 5 minutes at 80 rpm, RT. Cell counting was manually performed using Imaris (Bitplane) by an experimenter who was blinded to the conditions.

### Confocal imaging

10-15 z-stacks per animal were taken for the immunohistochemistry analysis. Around 50-100 z-stacks were taken for the dual-eGRASP density analysis, while 30-50 z-stacks of the image were used to analyse synaptic morphology. All samples were imaged using Leica SP8 confocal microscope or Leica Stellaris 5 with 63x objective with distilled water immersion and 10x objective lens was used for the immunohistochemistry. Images were taken within −2.9 to −3.8 mm bregma, spanning a range of ventral CA1 and -1.3 to 1.8 mm bregma for the BA.

Each dendrite expressing mScarlet-I or iRFP670 was identified and manually labelled as a filament using a filament tracer in IMARIS. To eliminate bias, other fluorescent signals were concealed. Cyan or yellow eGRASP signals were automatically detected and marked as cyan or yellow spheres. An overlap between cyan and yellow eGRASP signals was considered a yellow signal as it indicated that the presynaptic neurone of the synapse was c-Fos+ during memory formation. Dendrites lacking cyan or yellow eGRASP signals were excluded from further detailed analysis.

Spines on the designated mScarlet-I+ and iRFP670+ dendrites were manually reconstructed, and their diameter and volume were automatically detected. Each spine was categorised as either an activated spine or a non-activated spine based on the presence of a yellow or cyan eGRASP signal, determined through manual inspection. Measurements of spine head diameter, spine head volume, and spine length were performed using Imaris Filament Tracer. Importantly, the examiner conducting the spine 3D model reconstructions was kept blind to any eGRASP signals to prevent bias. Normalisation of the raw data was performed on a per-image basis to measure synaptic density and on a per-dendrite basis for morphology analysis. For the iRFP dendrite (for density analysis) or cyan-only spine morphology, the raw values were averaged, and then the raw values of each cyan-only spine and yellow spine were divided by this average. The resulting normalised values were utilised for subsequent statistical analysis.

### Statistical analysis

Data analysis was performed on Prism (GraphPad Software). The α value was set at 0.05 for all analyses.

---

**Fig S1.**
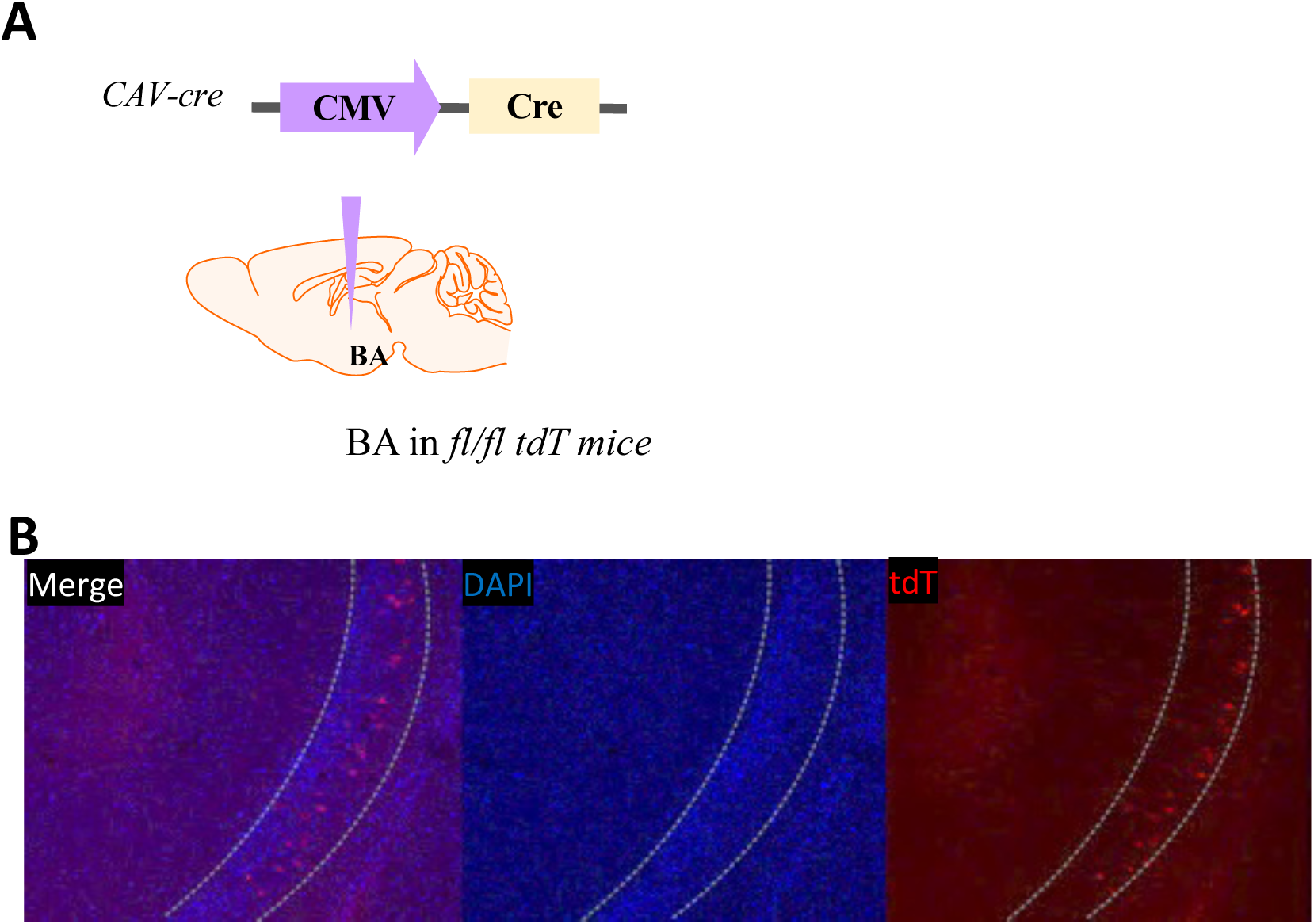
vCA1 neurones projecting to the BA are mainly residing in the superficial layer. (A) CAV-Cre virus injected into the BA of double floxed tdTomato mice (B) Images showing BA-projecting vCA1 neurones (tdT+)

**Fig S2.**
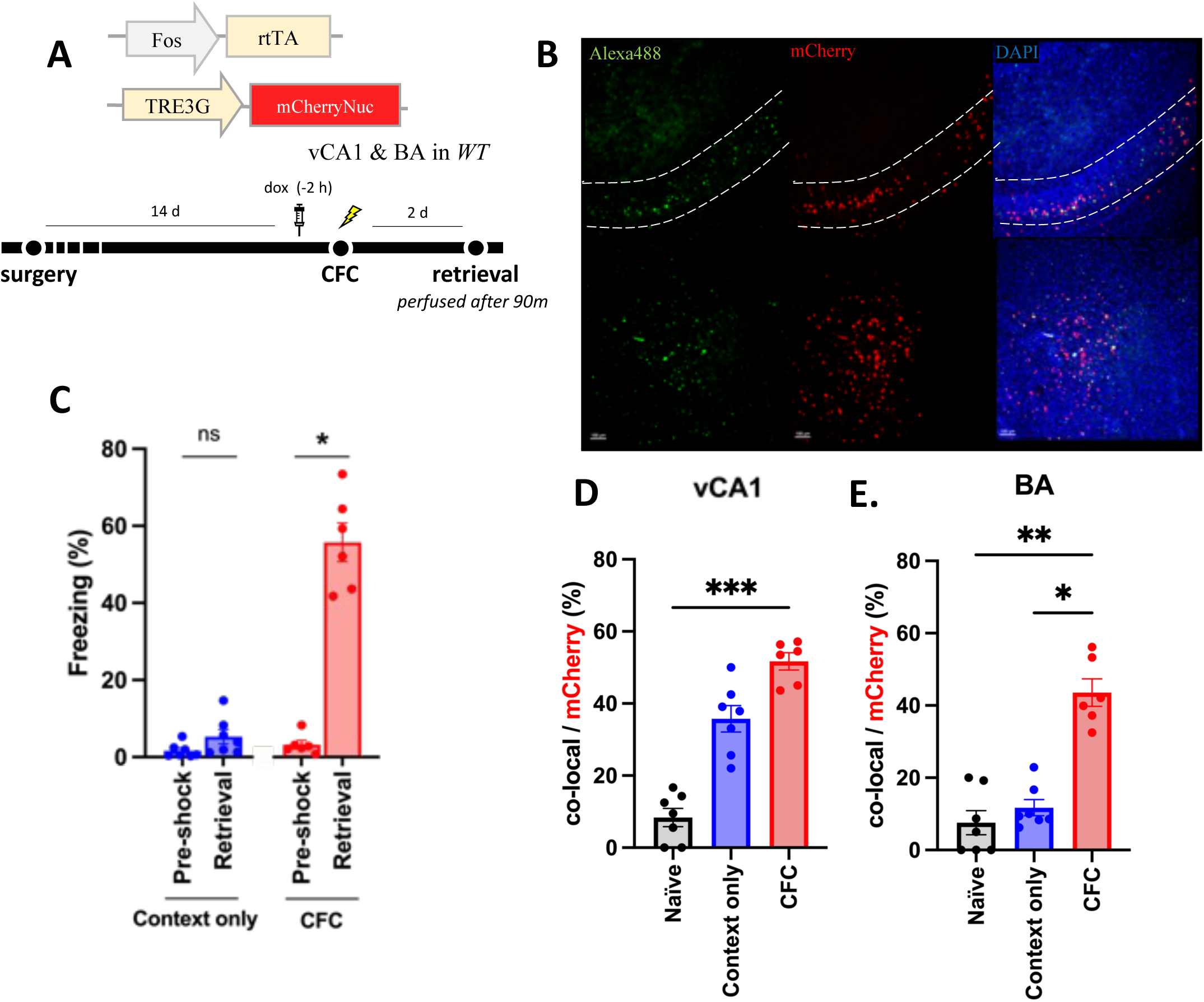
Engram reactivation of the vCA1 and BA during contextual fear conditioning. (A) AAV1/2 containing Fos-rtTA/ TRE-mCherryNuc was injected into the vCA1 and BA of wild type mice. (B) Representative image of the vCA1 (top) and BA (bottom). Neurons activated during the CFC were labelled in mCherry (middle) while neurons activated during memory retrieval were labelled in Alexa488 (left) (C) Freezing level comparing between to pre-shock session and memory retrieval session. Context only group, N = 7; CFC group, N = 6. Wilcoxon test. *P = 0.0312. Data are presented as mean ± SEM. (D-E) Reactivation ratio of engram cells in the vCA1 (**D)** and BA **(E**) in three groups: naïve, context only, and CFC. Naïve group, N = 7; context only group, N = 7; CFC group, N = 6. Wilcoxon test. ***P = 0.0002; **P = 0.0016; *P = 0.0256. Data are presented as mean ± SEM.

**Fig S3.**
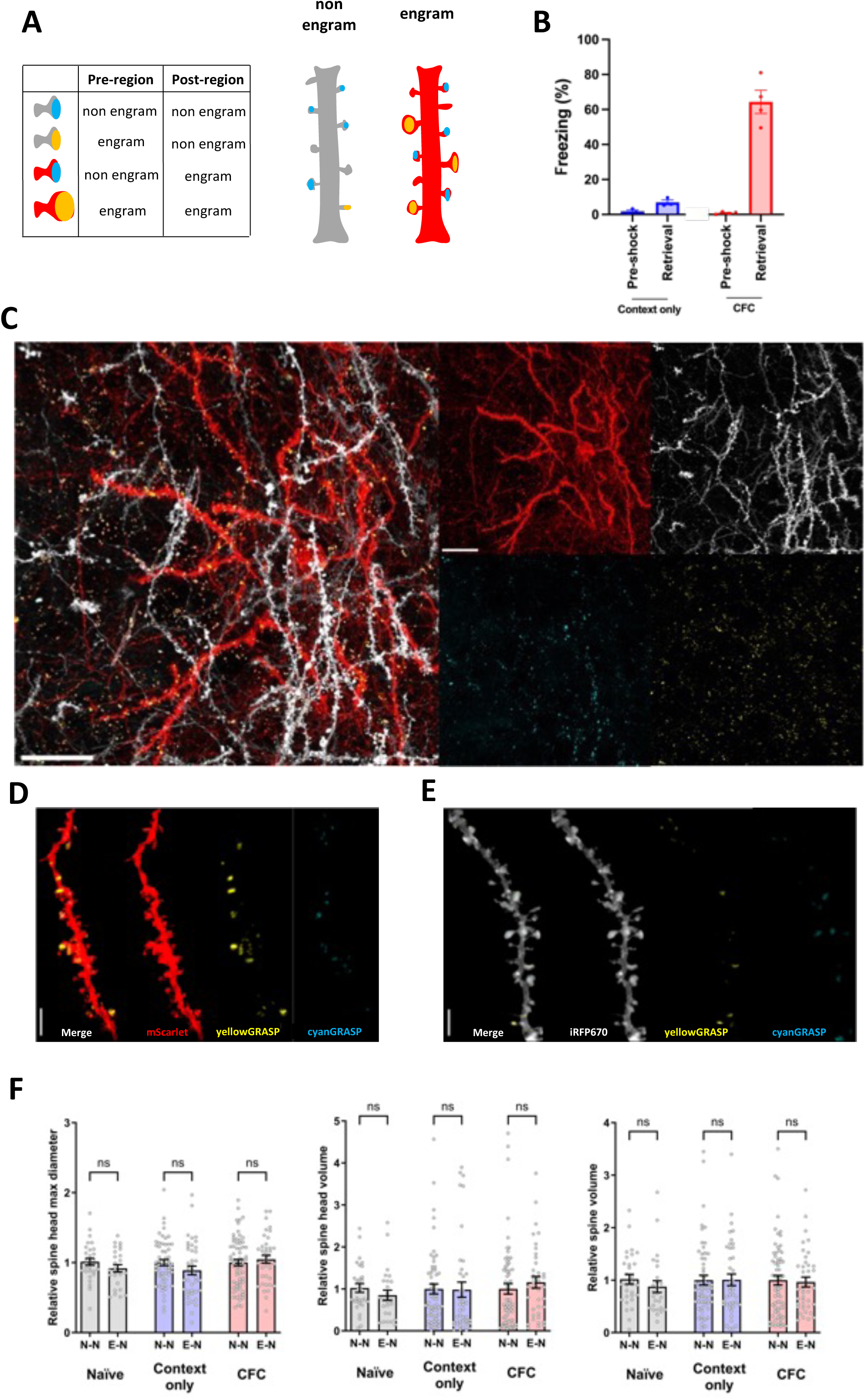
The overall synaptic connectivity from the vCA1 to the BA was not altered following the formation of memory. (A) Categorisation of the four synaptic populations, represented by distinct colours. (B) Freezing level during memory recall session for each group. Context only group, N = 3; CFC group, N = 4. Data are presented as mean ± SEM. (C) Illustrative images depicting both engram (E) and nonengram (N) dendrites featuring GRASP signals in the BA. Scale bar : 30 μm (D-E) Representative image of non-activated (D) and activated (E) GRASP signals displayed by each channel. Scale bar : 5 μm (F) Normalised spine head diameters, spine head volume and spine volume of dendrites from BA non engram cells. Sizes of the spines with yellow puncta were normalised to those of the spines with cyan-only puncta of the same dendrite. Each data point represents a spine. Naïve group N-N, *n* = 30; E-N, *n* = 26; Context only group N-N, *n* = 57; E-N, *n* = 41; CFC group N-N, *n* = 66; E-N, *n* = 39. Two-way ANOVA followed by Tukey’s multiple comparison test. Data are represented as mean ± SEM

**Fig S4.**
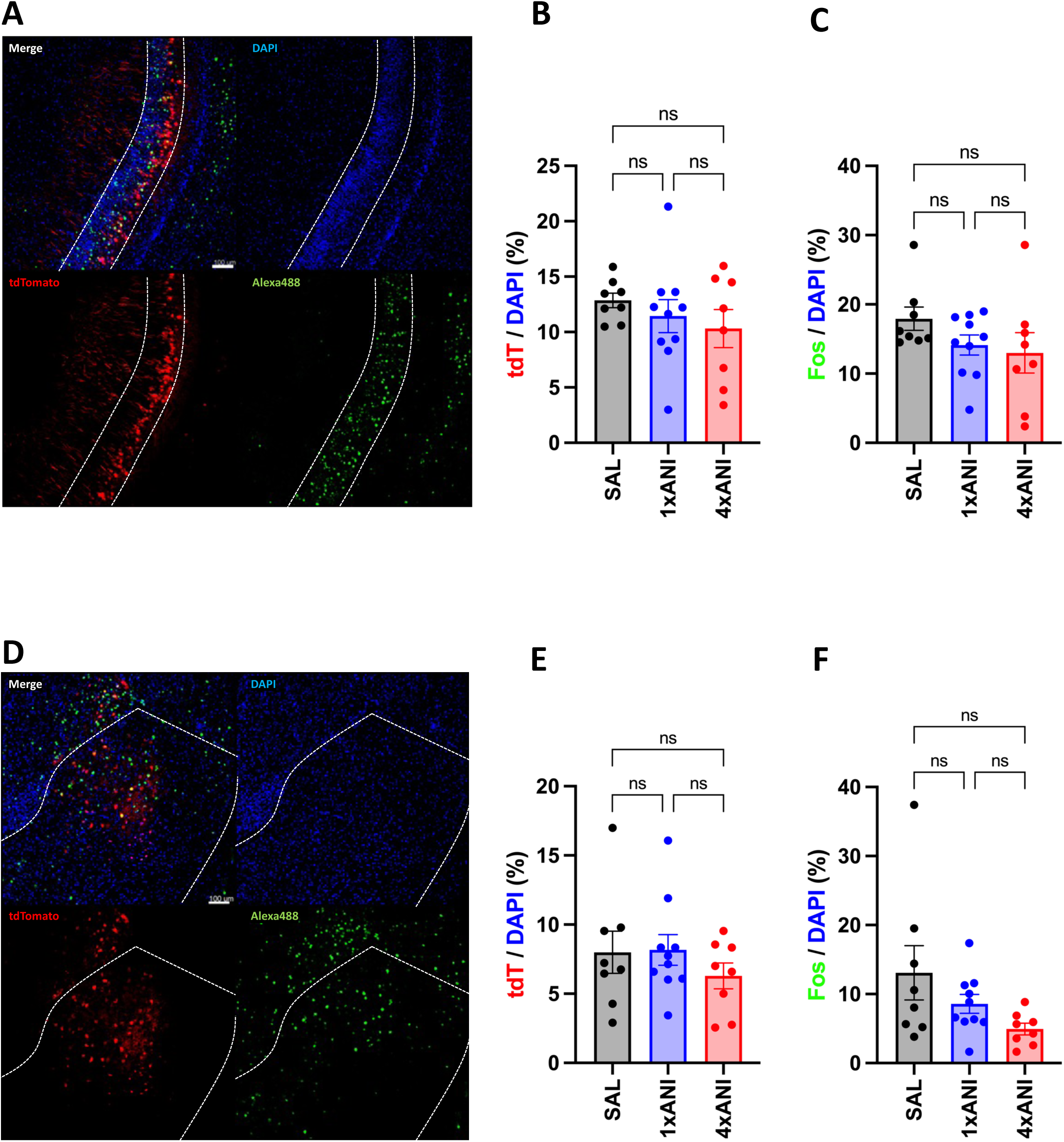
Engram labelling with PSI injections in Arc-TRAP mice. (A) Representative images of activated neurones during the fear conditioning (tdTomato+) in the vCA1 recall and neurones active during memory recall using cFos IHC (Alexa488). DAPI was stained to label the nuclei. (B-C) cFos IHC result for the vCA1. (D) Representative images of activated neurones during the fear conditioning (tdTomato+) in the BA recall and neurones active during memory recall using cFos IHC (Alexa488). DAPI was stained to label the nuclei. (D-E) cFos IHC result for the BA.

**Figure S5.**
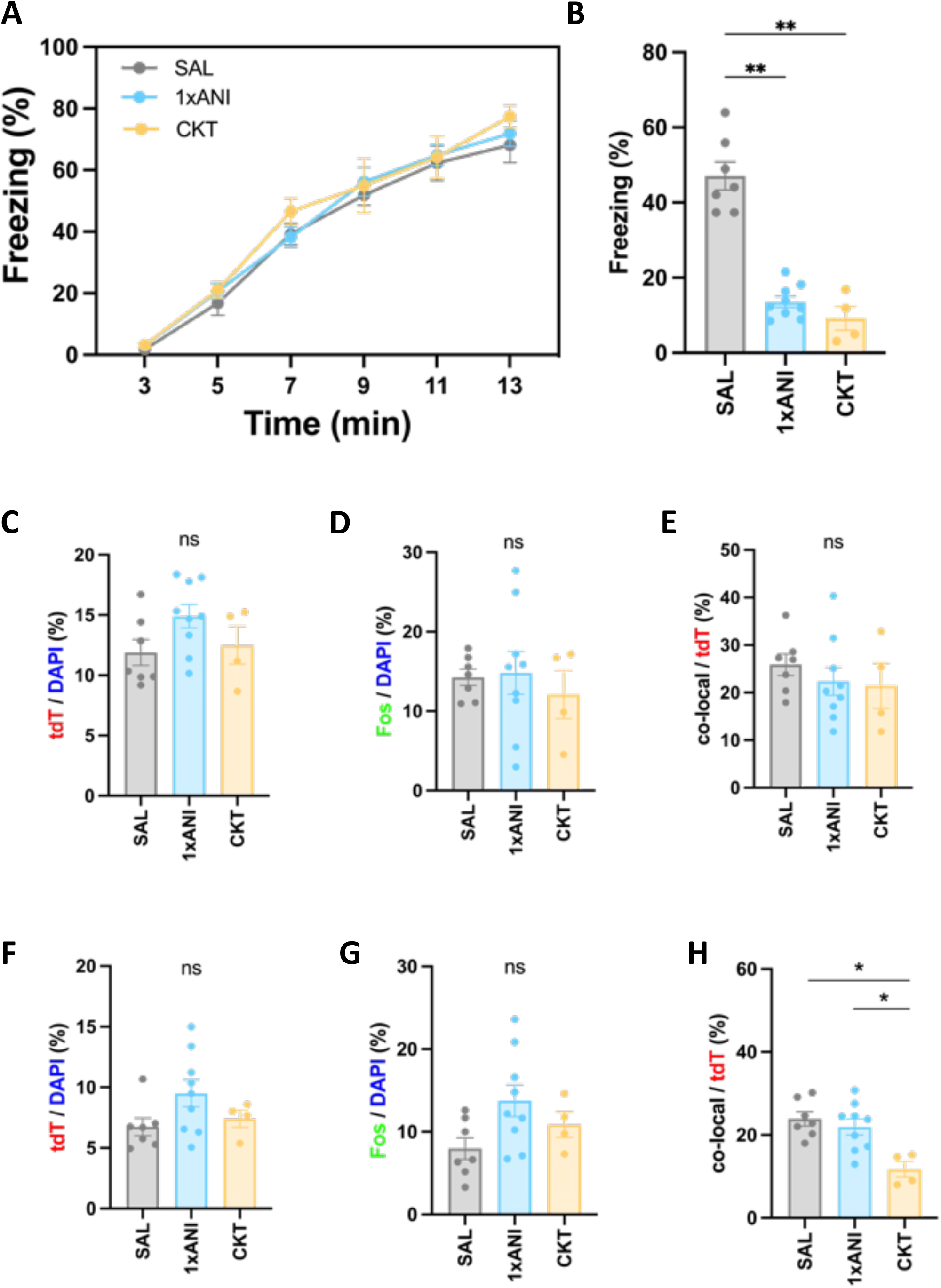
Anisomycin combined with cycloheximide induced stronger retrograde amnesia. (A) Acquisition plot for SAL, 1xANI, and CKT group during CFC. SAL group, N = 7; 1xANI group, N = 9; CKT group, N = 4. Data are presented as mean ± SEM. (B) Freezing level during memory recall. Kruskal-Wallis test followed by Dunn’s multiple comparison test. SAL vs. 1xANI, **P = 0.0067; SAL vs. CKT, **P = 0.0036. Data are presented as mean ± SEM. (C-E) Immunohistochemistry data of vCA1. Kruskal-Wallis test followed by Dunn’s multiple comparison test. Data are presented as mean ± SEM. (F-H) Immunohistochemistry data of BA. Kruskal-Wallis test followed by Dunn’s multiple comparison test. SAL vs. CKT, *P = 0.0148; 1xANI vs. CKT, *P = 0.0483. Data are presented as mean ± SEM.

**Fig S6.**
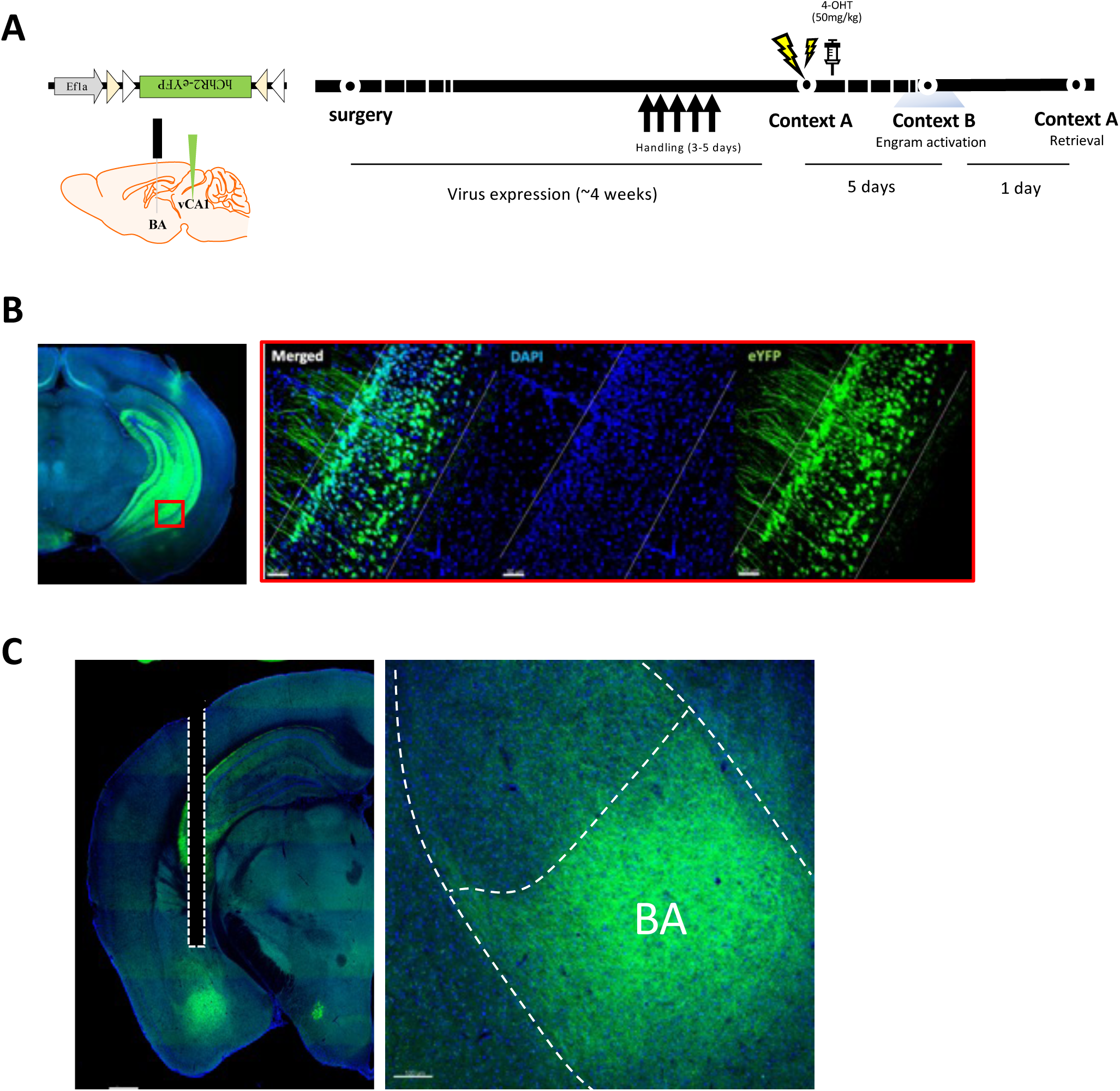
Representative images of virus expression for optogenetic behaviours. (**A**) Behavioural scheme (**B**) activated vCA1 neurones (**C**) vCA1 axonal projections to the BA expressed in ChR2-eYFP.

**Fig S7.**
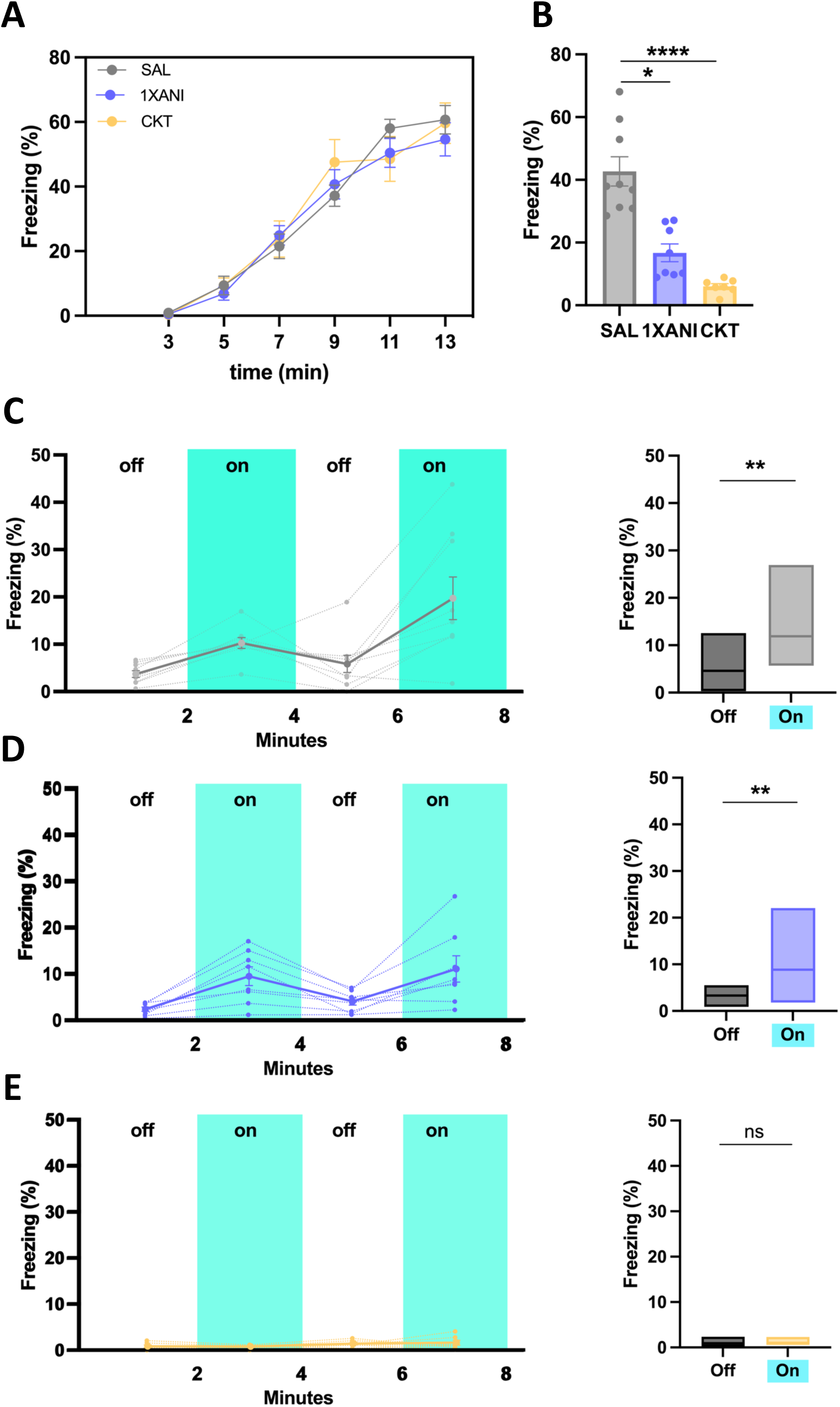
Memory cannot be artificially recalled when protein synthesis was blocked using combination of PSIs during memory acquisition. (A) Freezing level during memory acquisition. Data are presented as mean ± SEM. (B) Freezing level during contextual fear memory retrieval. SAL group, N = 9; 1xANI group, N = 8; CKT group, N = 7. Kruskal-Wallis test followed by Dunn’s multiple comparison test. *P = 0.0380; ****P < 0.0003. Data are presented as mean ± SEM. (C-E) Optogenetic activation of vCA1 engram axon terminals in BA of mice treated with different combination of PSI during memory acquisition. Mice expressing ChR2 in vCA1 engram cells show greater freezing during light-on epochs in saline-treated group (C) or by a single treatment of anisomycin (D) while absence of freezing behaviour was shown in mice treated with cocktail of PSI (E). Wilcoxon test. SAL group, **P = 0.0039; 1xANI group, **P = 0.0078. Data are presented as mean ± SEM (left) or median (right).

**Fig S8.**
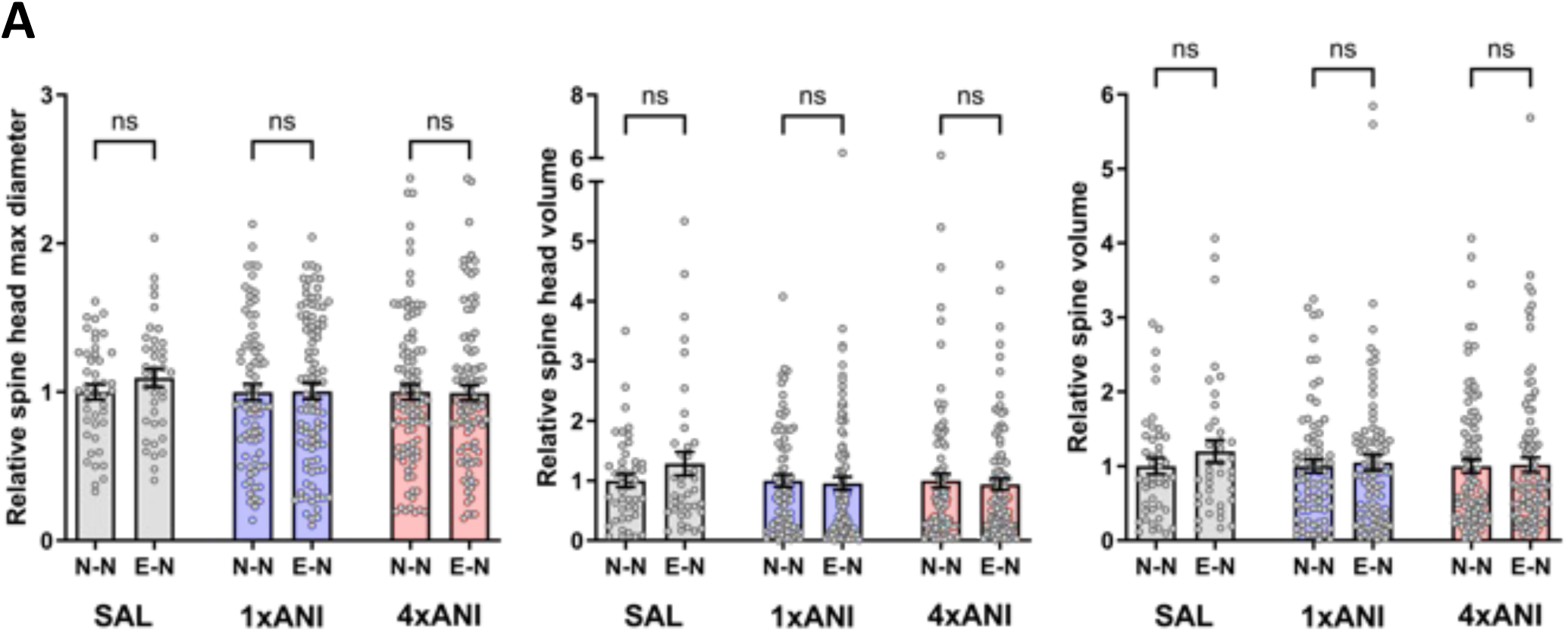
Normalised nonengram morphology after anisomycin treatment. (A) Normalised spine head diameter, spine head volume and spine volume of dendrites from BA nonengram cells. SAL group N-N, *n* = 44; E-N, *n* = 39; 1xANI group N-N, *n* = 77; E-N, *n* = 94; 4xANI group N-N, *n* = 96; E-N, *n* = 98. Two-way ANOVA followed by Tukey’s multiple comparison test. Data are presented as mean ± SEM.

